# Peripheral blood DNA methylation differences in twin pairs discordant for Alzheimer’s disease

**DOI:** 10.1101/468470

**Authors:** Mikko Konki, Maia Malonzo, Ida K. Karlsson, Noora Lindgren, Bishwa Ghimire, Johannes Smolander, Noora M. Scheinin, Miina Ollikainen, Asta Laiho, Laura L. Elo, Tapio Lönnberg, Matias Röyttä, Nancy L. Pedersen, Jaakko Kaprio, Harri Lähdesmäki, Juha Rinne, Riikka J. Lund

## Abstract

Alzheimer’s disease (AD) results from a neurodegenerative process that starts well before the diagnosis can be made. New prognostic or diagnostic markers enabling early intervention into the disease process would be highly valuable. As life style factors largely modulate the disease risk, we hypothesised that the disease associated DNA methylation signatures are detectable in the peripheral blood of discordant twin pairs. Reduced Representation Bisulfite Sequencing, single cell RNA-sequencing and gene array data were utilised to examine DNA methylation signatures and associated gene expression changes in blood and hippocampus, and targeted bisulfite sequencing in cross cohort validation. Our results reveal that discordant twin pairs have disease associated differences in their peripheral blood epigenomes. A subset of affected genes, e.g. *ADARB2* contain differentially methylated sites also in anterior hippocampus. The DNA methylation differences seem to influence gene expression in brain rather than in blood cells. The affected genes are associated with neuronal functions and pathologies. These DNA methylation signatures are valuable disease marker candidates and may provide insights into the molecular mechanisms of pathogenesis.

## Introduction

Alzheimer’s disease (AD) is an aging-associated neurodegenerative disorder and the most common cause of dementia. The molecular mechanisms of AD are not known in detail. The disease is characterised by accumulation of beta-amyloid plaques and hyperphosphorylated tau protein in the brain tissue. The pathological changes can start decades before the first clinical symptoms appear (Alzheimer’s Association. 2016,Querfurth and LaFerla. 2010,Dubois et al. 2016). Genetic factors contribute, however, do not fully determine the disease risk. In the early onset form of AD, which accounts for approximately 2-10 % of the cases, the symptoms can start already before age of 30 years. In 5-10 % of the early onset cases the disease is caused by autosomal dominant mutations in the genes *APP*, *PSEN1* and/or *PSEN2* (Alzheimer’s Association. 2016,Cuyvers and Sleegers. 2016). Majority of the AD cases are late onset form, which typically manifests after 65 years of age (Alzheimer’s Association. 2016,Cuyvers and Sleegers. 2016). Based on twin studies, the estimated heritability of the disease exceeds 50 % (Gatz et al. 2006). Genetic association with Apolipoprotein E (*APOE*) epsilon 4 (*ε4*) allele is common in both early and late onset form. Furthermore, genome-wide association studies have identified over 20 additional variants each contributing less than 1 % to the heritability of liability. Together with *APOEε4* these known variants explain approximately 29 % of the heritability (Lambert et al. 2013,Cuyvers and Sleegers. 2016) leaving a large fraction unexplained.

In addition to genetic factors, the morbidity of AD is modulated by modifiable lifestyle factors, such as physical activity and cognitive ability, nutrition, alcohol use, smoking and other illnesses (Reitz et al. 2011,Alzheimer’s Association. 2016). These factors may influence the disease process through epigenetic mechanisms, such as DNA methylation, by causing changes detectable in various tissues. Such changes may provide insights into the molecular mechanisms of the disease and carry the potential as informative biomarkers. The DNA methylation marks associated with AD in several brain regions have been discovered (Roubroeks et al. 2017,De Jager et al. 2014,Lunnon et al. 2014,Watson et al. 2016,Zhao et al. 2017). Whether similar marks exist in peripheral blood is still unclear (Fransquet et al. 2018). The question is complicated by the fact that methylation levels at CpG sites associated with the disease may be driven by genetic variants. Within-pair comparison of disease-discordant monozygotic (MZ) twin pairs with, in practice, identical genomes, the same age and sex, and who share many intrauterine and early life environmental factors provides a powerful approach for detection of the disease-associated methylation sites (Tsai and Bell. 2015). Dizygotic (DZ) disease-discordant twin pairs also provide improved sensitivity to the analyses as they share on average 50% of their genome and are of the same age and matched for prenatal and shared rearing environmental factors.

Modern technologies, such as Positron Emission Tomography (PET), enable monitoring of the disease status in the brains of living subjects, and are already utilised in diagnostics together with clinical examination and biomarkers to identify the individuals with first symptoms (Nordberg et al. 2010,Scheinin, Scheinin et al. 2011,Reitz et al. 2011). However, novel high-throughput methods enabling early detection and large scale monitoring of the disease status would be highly valuable. The aim of this study was to identify epigenetic marks associated with late onset AD in peripheral blood by comparing DNA methylation profiles of the disease-discordant twin pairs and to examine overlap in brain tissue. To evaluate potential as prognostic or diagnostic marker, one of the most interesting loci was selected for validation in extended twin cohorts from Finland and Sweden.

## Results

### Genetic risk load for Alzheimer’s disease in the Finnish study participants

To characterize the genetic risk load for AD in the Finnish study subjects we analysed the *APOE* genotypes and 21 loci previously associated with AD as a risk or protective variants (**Table 2**) (Lambert et al. 2013,Cuyvers and Sleegers. 2016). In this analysis, we included 9 full MZ and 12 full DZ AD-discordant twin pairs, 9 unrelated controls and 18 unrelated cases. Monozygotic co-twins were confirmed to have identical SNP profiles. Dizygotic co-twins were identical for the *DSG2*, *CASS4* and *SORL1* alleles, however, carried differences in 2-10 other variants. Most of the twin pairs (17 of 23 pairs) had *APOE* ε3ε3 genotype. To examine whether genetic risk load had an influence on the disease outcome, GRS were calculated for the study groups based on the 21 loci previously associated with AD (Table 2). As expected the monozygotic twin pairs did not differ for GRS. Also in dizygotic twin pairs, the GRS was not associated with the disease outcome.

### Blood DNA methylation marks associated with Alzheimer’s disease in the Finnish twin cohort

To examine whether epigenetic marks associated with AD can be identified in the peripheral blood of individuals with similar genetic risk profiles, we first identified the within-pair methylation differences associated with AD in 11 MZ twin pairs with RRBS as described in the Methods section. This comparison revealed 566 CpG sites with increased and 589 sites with decreased methylation associated with AD when using cut off values of at least 15 % average methylation difference and q-value ≤0.1 (Figure 1a, Table EV1a). Similarly, the CpG methylome differences were then identified in 12 DZ twin pairs discordant for AD. This comparison revealed 3340 sites with increased and 2201 sites with decreased methylation (Figure 1b, Table EV1b). We then examined the overlap of differentially methylated CpG sites detected in both MZ and DZ within-pair comparisons with 100 bp genomic window. This revealed 74 common sites with consistent methylation difference present in both zygosity groups and with a combined q-value ≤0.02 (Figure 1c-d, Table EV1c). After merging the CpG sites within 100 bp distance from each other into regions, we found 39 genomic regions containing differentially methylated sites within the twin pairs discordant for AD. The most significant regions were closest to, or within genes, such as *TSNARE1, DEAF1, ADARB2.* In addition to autosomes, we found one additional differentially methylated site in chromosome X upstream (-48,704 bp) from the gene *AR*. No methylation differences were detected in the mitochondrial DNA. In a summary, DNA methylation changes in at least 39 genomic regions in autosomes and one in chromosome X were found to be associated with AD in peripheral blood of AD-discordant Finnish twin pairs. These regions are henceforth referred to as AD-associated loci in blood.

**Figure 1.**
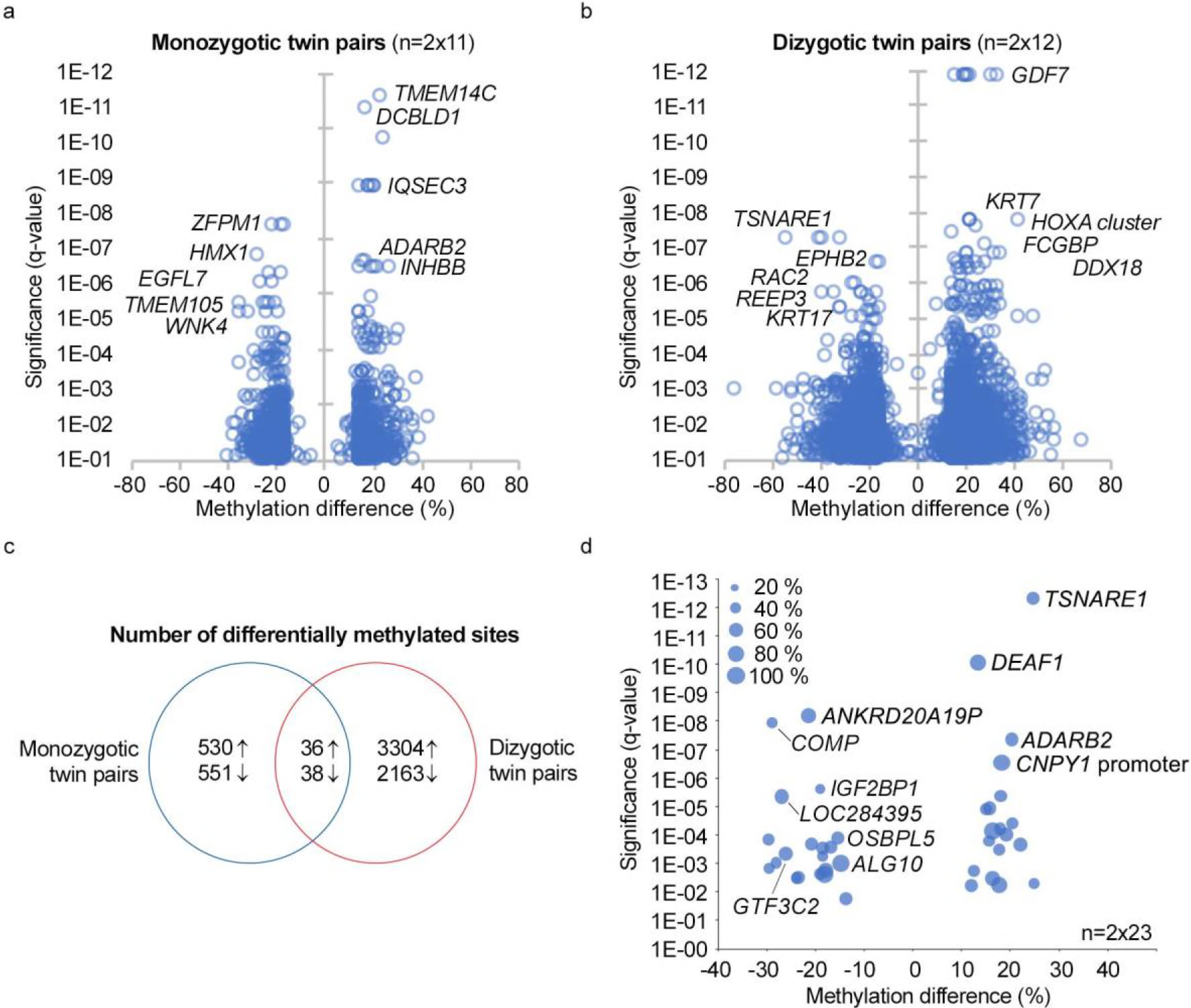
Peripheral blood CpG methylation differences in Finnish twin pairs discordant for Alzheimer’s disease. Peripheral blood CpG methylomes of 11 monozygotic (MZ) and 12 dizygotic (DZ) Finnish twin pairs discordant for Alzheimer’s disease (AD) were profiled with Reduced Representation Bisulfite Sequencing. Differentially methylated sites associated with AD in autosomes were identified using RADMeth algorithm and ±15 % average methylation difference and q-value cut off 0.1 in a) MZ and b) DZ twin pairs separately. c) Differentially methylated sites within 100 bp distance in both MZ and DZ twin pairs were extracted to identify overlaps between the AD-associated sites detected in both groups. d) The AD-associated autosomal sites within 100 bp distance from each other were merged into 39 regions (Table EV1). Combined average methylation difference and q-value distribution of these 39 regions associated with AD in both MZ and DZ twin pairs is illustrated in the Figure. The size of the dot indicates the percentage of the twin pairs, which had the required 10x coverage in the region. The closest or overlapping gene is indicated for the selected differentially methylated regions.

### DNA methylation changes associated with Alzheimer’s disease in anterior hippocampus and overlap with peripheral blood

To examine whether AD-associated loci in blood were also affected in the brain region crucial for memory formation and learning, DNA methylation was profiled with the RRBS in the anterior hippocampus from postmortem tissue sections collected from six cases with AD and six controls (Table 1). Two cases were excluded from the final analysis as one was identified as an outlier in the principal component analysis and according to the neuropathological examination one sample was from amygdala. Comparison of cases and controls revealed increased methylation at 114 sites and decreased methylation at 87 genomic sites (at least 15 % methylation difference, q-value ≤0.05), overlapping or closest to the 176 gene identifiers, in the anterior hippocampus (Figure 2a). These sites are henceforth referred to as AD associated loci in brain. Comparison of the gene identifiers closest to the AD associated loci in brain to those detected in blood revealed four overlapping genes, *ADARB2, OLFM1, CTDP1* and *CAMKMT/SIX3* (Figure 2b). In the gene *ADARB2,* the differentially methylated sites in the blood and hippocampus were localised within the same region chr10:1404752-1405717 in exon 3, in the proximity of the exon 3 - intron junction.

**Figure 2.**
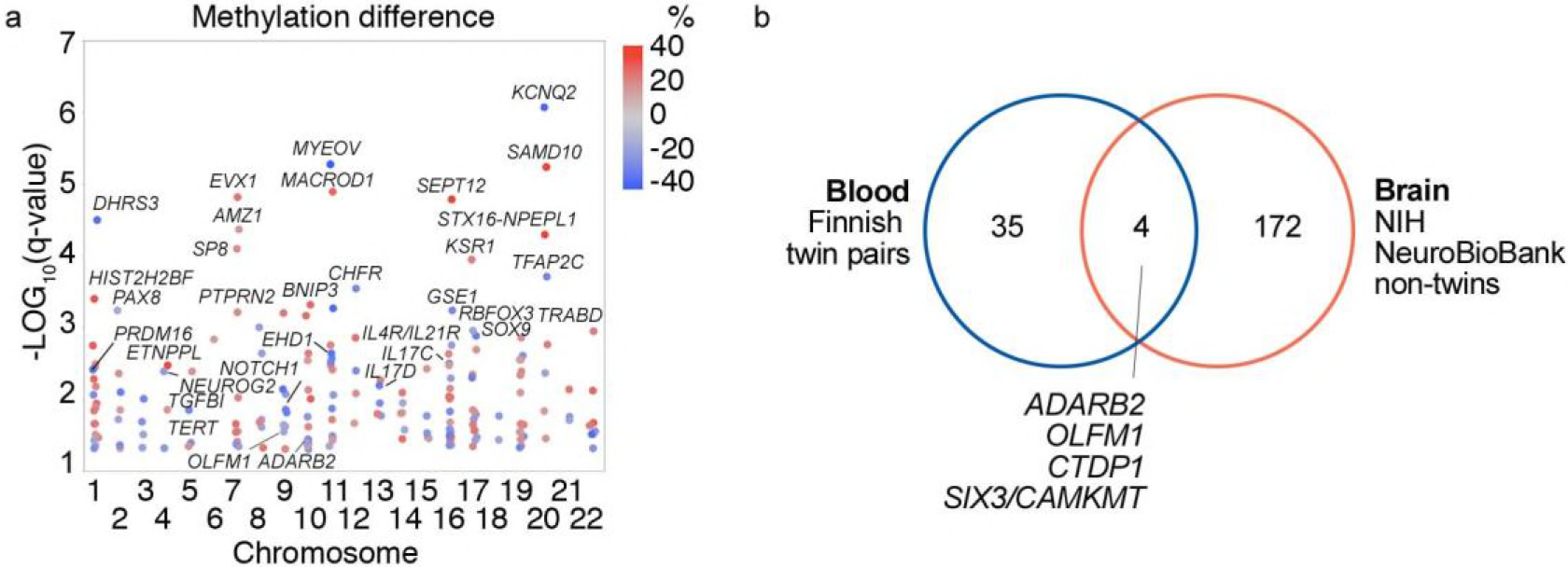
Differentially methylated sites associated with Alzheimer’s disease in the anterior hippocampus. DNA methylation profiles of anterior hippocampus samples of 10 non-twin subjects, including six references and four with Alzheimer’s disease (AD), were examined with Reduced Representation Bisulfite Sequencing. The samples were from NIH NeuroBioBank. **a**) Differentially methylated sites (201 CpGs) were identified with MethylKit package in R by using a minimum of 15 % methylation difference and q-value below 0.05 as filtering cut-off (Table EV1). The 201 differentially methylated CpG sites overlapped or were closest to 176 genes (AD-associated genes in brain). Examples of the genes are shown in Figure. **b**) Overlap of genes with AD-associated DNA methylation marks in both peripheral blood and anterior hippocampus of brain.

**Table 1.** Characteristics of the samples utilized in the study.

| Study and sample type | Controls |  | Cases |  |
| --- | --- | --- | --- | --- |
|  | No of subjects (male/female) | Age range (years) | No of subjects (male/female) | Age range (years) |
| **Finnish cohort (n=97)** |  |  |  |  |
| *Peripheral blood (n=93)* |  |  |  |  |
| ADD<sup>a</sup> MZ twin pairs | 12/9 | 67-86 | 12/9 | 67-86 |
| ADD DZ twin pairs | 6/6 | 70-72 | 6/6 | 70-72 |
| Unrelated individuals | 1/8 | 61-77 | 9/9 | 70-86 |
| *Peripheral blood mononuclear cells (n=4)* |  |  |  |  |
| MZ twin pairs discordant for episodic memory | 2/0 | 76 | 2/0 | 76 |
| **Swedish Cohort (n=298)** |  |  |  |  |
| *Peripheral blood (n=298)* |  |  |  |  |
| ADD MZ twin pairs | 5/5 | 62-83 | 5/5 | 63-83 |
| ADD DZ twin pairs | 11/8 | 63-84 | 11/8 | 63-84 |
| Incident<sup>b</sup> ADD MZ twin pairs | 15/23 | 62-85 | 15/23 | 62-85 |
| Incident ADD DZ twin pairs | 33/49 | 53-88 | 33/49 | 53-88 |
| **NIH NeuroBioBank (n=12)** |  |  |  |  |
| *Anterior hippocampus with entorhinal cortex (n=12)* |  |  |  |  |
| Unrelated individuals | 4/2 | 56-89 | 4/2 | 57-90 |
| **Total (n=407)** | **89/110** | **53-90** | **97/111** | **53-90** |

<sup>a</sup>Alzheimer's disease-discordant.<sup>b</sup>Sample collected 0.5-18.5 years before disease onset.

### Gene expression levels in hippocampus, whole blood and individual lymphocytes

Expression of the 40 genes overlapping or closest to the AD-associated loci in blood (Figure 1d, Table EV1c) were examined in the data from hippocampi (GDS810) including controls (n=9), incipient (n=7), moderate (n=8) and severe (n=7) AD cases (Blalock et al. 2004) and whole blood (GSE63061, Europeans) of 122 controls and 121 cases with AD (Sood et al. 2015) available at GEO database. In the hippocampus expression of 20 genes and several transcripts from the *PCDHG* cluster was detected. The most significant difference was increased expression of the *ADARB2* gene in patients with severe AD in comparison to controls (1.85-fold, p=0.005, adj. p=0.22, Table EV2, Figure EV1). Furthermore, a trend of differential expression, including genes *ARAP2*, *OLFM1*, *CDH11*, *CLIP2, COMP, CTDP1*, *IRX4* and transcripts from *PCDHG* cluster, was detected in at least one of the patient groups in comparison to controls (nominal p≤0.05, Table EV2, Figure EV1). In addition, expression levels of the genes *OLFM1*, *DEAF1* and *GTF3C2* were reported to correlate with either MiniMental Status Examination or neurofibrillary tangle scores in the original study (Blalock et al. 2004). No significant differences were detected for the other genes. HG133A array used in the original study (Blalock et al. 2004) does not contain probes for *TSNARE1* and *SORCS2* genes. In blood, expression of 20 genes and transcripts from PCDHG cluster were detected, however, there were no differences between the controls and cases with AD (Table EV2b, Figure EV2). Expression of these genes was further examined in individual peripheral blood lymphocytes of twin pairs discordant for episodic memory and the other pair also for mild AD. The proportions of different blood cell types expressing these genes were similar between the co-twins (Figure EV3) and no major differences were detected in the expression of the genes within the twin pairs (Figure EV4). In conclusion, several of the genes closest to the AD-associated differentially methylated loci in blood seem to be differentially expressed in the hippocampi of individuals with AD, including *ADARB2*, *OLFM1* and *CTDP1,* methylation of which was affected in both blood and brain. Analysis of blood did not provide evidence that the genes would display major disease associated functional significance in the PBMC populations.

### Targeted validation of disease-associated methylation changes in *ADARB2* gene in Finnish and Swedish twin pairs discordant for AD

The CpG methylation of the region (chr10:1,405,405-366) in *ADARB2* gene was analysed with targeted bisulfite pyrosequencing in 62 of AD discordant twin pairs including independent 29 Swedish twin pairs. Association of the CpG methylation with AD was analysed using linear mixed effects model (lme) (Bates et al. 2015) including disease status, gender, zygosity, age and their interactions as fixed effects, and twin pairs nested with genomic position and country of origin as random effects. Influence of *APOE* genotype (*ε34/ε44* and *ε33*) was examined separately, including disease status and gender as fixed effects, to ensure sufficient number of observations in each subgroup. According to our results, influence of country of origin on the CpG methylation level was not significant (ΔAIC 2.0, ANOVA p-value 1). Increased CpG methylation level of the region chr10:1,405,405-366 was validated to be associated with the AD (Wald t-value 4.68) and was influenced by interaction with gender, zygosity and age. Separate examination by sex revealed that the CpG methylation level was increased in male cases (Wald t-value 6.10) (Figure 3a, Figure EV5). The methylation difference of the region was higher in DZ than MZ twin pairs (estimate 7.67, standard error (SE) 1.44, Wald t-value 5.34) (Figure 3b). In addition, in males the CpG methylation difference between discordant twin pairs increased with age (estimate 1.00, SE 0.15, t-value 6.43) (Figure 3c). Interestingly, in females the CpG methylation was increased in cases with APOE *ε*34/*ε*44 genotype (Wald t-value 3.44), however, not in cases with *ε*33 genotype (Wald t-value -1.67) (Figure 3d). In male cases the CpG methylation level was higher in both *APOE* genotype groups (Wald t-value >3.83), and in cases the level in *APOE* ε34/ε44 group was higher than in *ε*33 genotype group (Wald t-value 4.59). Of note, visual examination revealed that also the male cases with *ε*33 genotype had polarized into high and low/intermediate CpG methylation level groups (Figure 3e). In conclusion, our findings from targeted analysis confirm the association of CpG methylation in region chr10:1,405,405-366 with AD and the methylation is influenced by gender, age, zygosity and *APOE* genotype.

**Figure 3.**
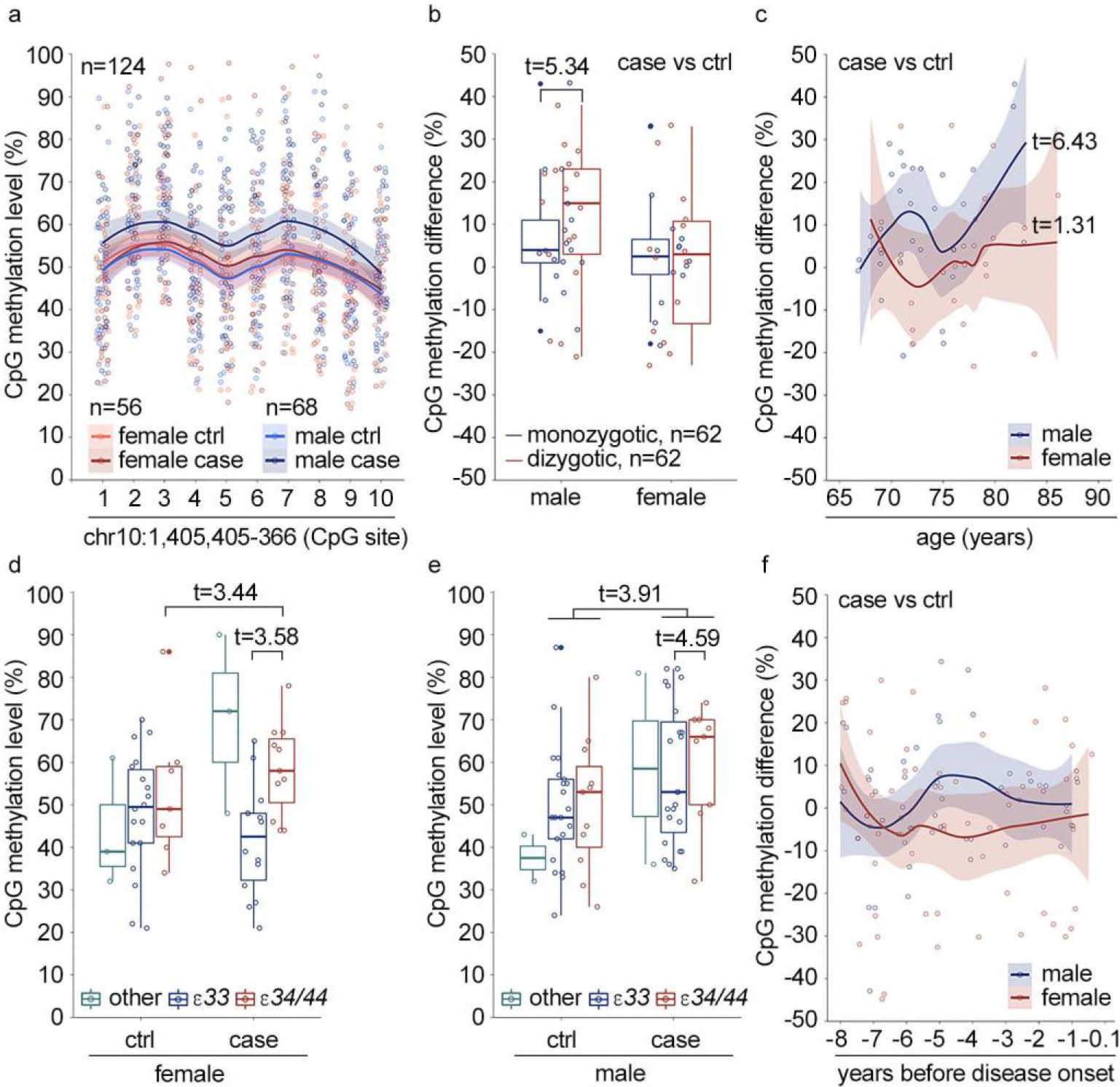
The DNA methylation level of region chr10:1,405,405-366 in *ADARB2* gene is associated with Alzheimer’s disease after disease onset and is influenced by gender, zygosity, age and *APOE* genotype. The DNA methylation level of region chr10:1,405,405-366 in exon 3 of *ADARB2* gene was measured with targeted pyrosequencing in blood DNA samples collected from Finnish (33 pairs) and Swedish (29 pairs) twin pairs discordant for Alzheimer’s disease (AD). The association of the disease status with DNA methylation was examined with linear mixed effects model including gender, zygosity, age and interaction terms as fixed effects and twin pair information nested with genomic position as a random effect. The country of origin was not associated with the CpG methylation level and was excluded from the final models. **a**) The CpG methylation level in male and female controls (ctrl) and cases. Wald t-value for disease association including both genders is 4.68, for males 6.10 and for females 1.17. **b**) Influence of zygosity of CpG methylation difference between twin pairs discordant for AD. **c**) Influence of age on the CpG methylation difference between AD discordant twin pairs. Influence of *APOE* genotype on the CpG methylation level was examined separately including only gender as a covariate. Only group ε33 and combined groups ε34/44 were included in the statistical analysis. The CpG methylation levels for **d**) female and **e**) male ctrls and cases in different APOE groups. **f**) CpG methylation difference in AD discordant twin pairs before onset of the diseases (not predictive for disease outcome based on cox mixed effects model). In the Figure3b-f the representative data for CpG site 6 in the region is shown, and in the Fig.3a, c and f the lines are the smoothed conditional means with 0.95 confidence intervals area.

### CpG methylation status of region chr10:1,405,405-366 in *ADARB2* gene does not predict the disease outcome

To determine whether CpG methylation level in region chr10:1,405,405-366 predicts the disease development we used Cox mixed effects model and examined association of CpG methylation with disease outcome in samples collected 0.5-18.5 years before disease onset from 120 discordant twin pairs from Sweden. The gender and age at blood draw were included as covariates in the analysis. According to the results the disease outcome was not associated with the DNA methylation level changes of the region before disease onset (Figure 3f).

## Discussion

Our results show that DNA methylation differences associated with AD can be detected in the peripheral blood in at least 40 genomic regions. Although functional importance of the affected regions in transcriptional regulation requires further studies, our results suggest that the expression of the closest genes is affected in brain rather than in peripheral blood. Consistently several of these genes have been previously linked to neurological functions or pathologies. Among the most interesting findings was the AD-associated methylation signature in the exon 3 of *ADARB2* gene, which was detected in both peripheral blood and anterior hippocampus in brain tissue. Furthermore, transcription of *ADARB2* was associated with AD in hippocampus. Based on previous studies, expression of *ADARB2* is brain specific, however, its function has not been fully characterized. *ADARB2* may have a function in the inhibition of RNA editing by ADAR1 and ADAR2 proteins (Tan et al. 2017,Oakes et al. 2017,Gaisler-Salomon et al. 2014). Importantly, a recent study showed that mice lacking the corresponding exon 3 of *ADARB2* gene display impaired hippocampus-dependent memory formation, learning and regulation of genes implicated in synaptic functions (Mladenova et al. 2018). Consistently, a variant in *ADARB2* gene has been associated with accelerated cognitive decline after conversion from mild cognitive impairment (MCI) to AD (Lee et al. 2017). With targeted validation including independent twin pairs discordant for AD from both Finnish and Swedish cohorts, we further found that the methylation level in *ADARB2* was influenced by *APOE* genotype, age and gender, which all also influence risk of AD. Although these results together suggest potential importance of *ADARB2* in the molecular pathology of AD, the high inter-individual variance and relatively small within-pair differences in blood suggest that methylation at this region alone is unlikely to have value as a diagnostic marker. Furthermore, CpG methylation in *ADARB2* was not predictive for AD when measured before disease onset.

Similarly to *ADARB2,* most of the other differentially methylated regions in blood overlapped or were closest to genes that are highly expressed in brain or have been previously linked to neuronal functions or pathologies, such as synaptic function, tau levels, amyloid deposition, cognition, memory formation and schizophrenia (Nakaya et al. 2012,Bertelsen et al. 2015,Vulto-van Silfhout et al. 2014,Bartelt-Kirbach et al. 2010,Deming et al. 2017,Fanous et al. 2012,Goes et al. 2015,Li et al. 2015,Mladenova et al. 2018,Reitz et al. 2013,Duan et al. 2014). For example, the most significant difference was detected in *TSNARE1* gene, that has been previously associated with rate of cognitive decline in late MCI (Li et al. 2015,Gu et al. 2015,Sleiman et al. 2013). Another example is *OLFM1,* highly expressed in brain, is implicated in neurogenesis and regulation of neuronal functions (Anholt. 2014,Nakaya et al. 2012,Fagerberg et al. 2014, Nakaya et al. 2013,Bertelsen et al. 2015). The functional importance of these genes remains to be further elucidated. In particular, further studies are needed to elucidate whether combinations of differentially methylated regions in multiple AD-associated loci would increase sensitivity in distinguishing AD cases from cognitively preserved controls and to evaluate the value of these changes as predictive disease markers.

## Materials and Methods

### Study participants and samples

A total of 395 peripheral blood samples from Finnish and Swedish cohorts and 12 brain samples from NIH NeuroBioBank, were analysed in this study (**Table 1**). In more detail, the older Finnish twin cohort (Kaprio and Koskenvuo. 2002,Kaprio. 2013) of 2,483 individuals, born 1922-1937, had been previously screened for cognitive functions by phone interview (Jarvenpaa et al. 2002) and the discordant twin pairs were invited to neuropsychological testing, brain Magnetic Resonance Imaging (MRI), Positron Emission Tomography (PET) (Jarvenpaa et al. 2003,Jarvenpaa et al. 2004,Scheinin, Aalto et al. 2011,Virta et al. 2008,Scheinin, Scheinin et al. 2011) and collection of EDTA blood samples. Based on the overall evaluation a diagnosis of AD was made (29 pairs). In addition to the original cohort, four twin pairs born in 1915-1950 were included in the study. Furthermore, BD Vacutainer^®^ CPT™ samples were collected two MZ twin pairs (age 76 years) discordant for episodic memory, the other pair also discordant for mild AD. These two twin pairs were from a younger Finnish twin cohort (born in 1938-1944). The blood DNA samples from Swedish twin pairs (149 pairs), born 1907-1953, were from the Swedish Twin Registry (Magnusson et al. 2013) and had been collected before (120 pairs) or after (29 pairs) the onset of the disease. Participants originated from three sub-studies of aging within the Swedish Twin Registry (Magnusson et al. 2013), namely the Swedish Adoption/Twin Study of Aging (SATSA) (Finkel et al. 2005), the Study of Dementia in Swedish Twins (HARMONY) (Gatz et al. 2005), and TwinGene (Magnusson et al. 2013). Disease information was available through linkage to nationwide health registers for all Swedish twins, and in addition AD was clinically evaluated as part of the SATSA and HARMONY study (Gatz et al. 1997). Furthermore, our study material included fresh frozen post-mortem brain tissue from anterior hippocampus, including dentate gyrus and entorhinal cortex. The samples had been collected from 12 adults (57-90 years), including six patients with AD and six non-demented controls. Five of the individuals were white and seven had unknown ethnic origin.

### Nucleic acid extractions

The DNA was extracted from EDTA blood with Qiagen QIAamp DNA Blood kit and from fresh frozen tissue with QIAamp DNA Micro kit. Qubit 3.0 dsDNA HS Assay, Thermo Scientific NanoDrop 2000 and Fragment Analyzer HS Genomic DNA assay (Advanced Analytical) were used for the quality analysis and quantitation.

### Targeted genotyping and variant data analysis

Genotyping of *APOE* alleles and 21 SNPs previously associated with AD (Lambert et al. 2013) (Table 2) was performed with Illumina TruSeq Custom Amplicon assay and sequencing (500 cycles) with Illumina MiSeq. Illumina BaseSpace was used for variant analysis. Genetic risk scores (GRS) were calculated based on 21 genetic variants by summing up the number of risk or protective alleles, weighted by the effect size from the stage 1 and 2 meta-analysis (Lambert et al. 2013). Influence of GRS on the disease outcome was examined with generalized linear model in R version 3.4.3 (R Core Team. 2017). The influence of sex, age and *APOE* genotype on the model was examined separately due to small number of samples. Packages lmtest (Zeileis and Hothorn. 2002), multiwayvcov (Graham et al. 2018) and fmsb (Nakazawa. 2017) were utilized in the model and calculation of Nagelkerke’s R square.

**Table 2.**
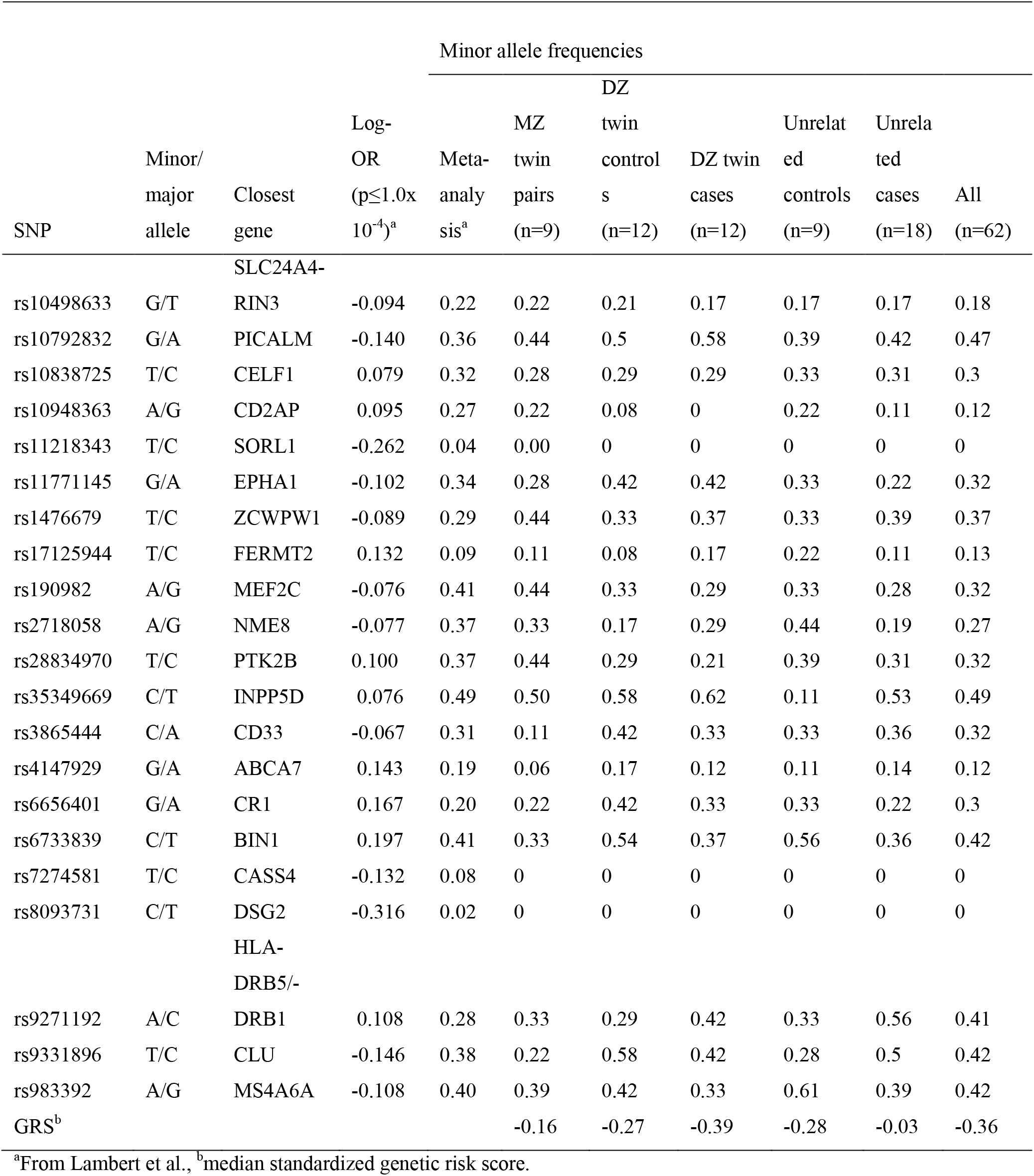
Genetic risk load for Alzheimer’s disease in the Finnish twin pairs and unrelated individuals.

### Reduced representation bisulfite sequencing and data analysis

The libraries for RRBS were prepared as previously described (Boyle et al. 2012,Konki et al. 2016) and were sequenced (1x50bp) with Illumina HiSeq2500/3000. The trimmed reads (Trim Galore v0.4.1 (Krueger. 2015) were mapped to hg19 (blood) or hg38 (brain) reference genome using Bismark v0.14.5/0.15.0 (Krueger and Andrews. 2011). The differentially methylated sites in blood were identified with RADMeth (Dolzhenko and Smith. 2014) using CpG sites with coverage ≥ 10x in both twins in a pair, and at least four pairs per study group, as an input. The differentially methylated regions were filtered according to combined p-value adjusted for multiple testing (< 0.1) followed by absolute average within-pair methylation difference (≥ 15%), and central interval (70% interval < 0 or 70% interval > 0) for these sites. Also for brain samples the CpG sites with coverage ≥10x and less than 99.9th percentile of coverage in each sample were included. Methylkit v.1.1.7 (Akalin et al. 2012) was used to detect differentially methylated CpG sites with a minimum and consistent difference of 15 % and q-value ≤0.05 as a cut-off. Bisulphite conversion efficiency of all the samples was above 99 % according to the lambda spike-in control.

### Targeted bisulfite pyrosequencing analysis

The PCR primers: 5’-gtaatttagtggtgttgttgaat-3’, 5’-biotin-cctaacccccaaccaacttcttactac-3’; and the sequencing primer: 5’-gggttgagttaagtgtgtttggtaga-3’ for the region chr10:1405336-1405409 (hg19), were designed with Qiagen PyroMark AssayDesign SW 2.0. The samples were prepared with Qiagen EpiTect Fast Bisulfite Conversion and Qiagen PyroMark PCR kits and sequenced with Qiagen PyroMark Q24 Advanced system. The amplicons were analysed with PyroMark Q24 Advanced software version 3.0.0. The statistical analysis was carried in R v3.4.3 (R Core Team. 2017) using packages lme4 v1.1-15 (Bates et al. 2015), car v2.1-6 (Fox and Weisberg. 2011), survival 2.42-4 (Therneau. 2018b), coxme 2.2-10 (Therneau. 2018a) and ggplot2 v2.2.1 (Wickham. 2009). Association of AD with CpG methylation was examined with lme4 package. Influence of covariates and their interaction in the model was examined, including gender, age, zygosity and *APOE* genotype as fixed effects and twin pair nested with genomic position and country of origin as random effects. The cox mixed-effects model with coxme package was utilized to examine CpG methylation level as a prognostic marker of disease outcome. In this model only age and gender were included as fixed effects and twin pair information nested with genomic position as a random effect.

### Single cell transcriptome analysis

PBMCs were isolated from the BD Vacutainer^®^ CPT™ Cell Preparation tubes. Single cell RNA-sequencing (scRNA-seq) libraries were prepared using Chromium™ controller, Single Cell 3’ Reagents kit (10X Genomics), targeting at the recovery of 3000 cells per sample, and were sequenced with Hiseq2500. The data was preprocessed using Cell Ranger v. 1.2.0 (10x Genomics) and the GRCh38 genome reference. The filtered gene-barcode unique molecular identifier (UMI) count matrix of the aggregated sample (Cell Ranger aggr tool) was normalized using a global-scaling normalization from the Seurat R package v. 1.4.0.9 (Butler et al. 2018). Finally, the full Seurat scRNA-seq analysis was performed for each sample individually. Cells with unique gene count over 1,750 or proportion of mitochondrial genes over 10% were first filtered out. Ten most significant principal components were selected for the graph-based clustering and the different PBMC cell types were identified using canonical markers and the cell type frequencies were estimated. Paired t-test was used to test differential expression between sample groups.

### Public data used

The data from GDS810 (Blalock et al. 2004) and GSE63061 (Sood et al. 2015) available at NCBI GEO database was analysed with GEO2R using default settings (Edgar et al. 2002,Barrett et al. 2013).

## Acknowledgements

We acknowledge Finnish Functional Genomics Centre, Biocenter Finland, University of Turku and Åbo Akademi University for the infrastructure support, NIH NeuroBioBank for the brain samples and Sinikka Collanus for the H&E staining of the brain samples. This study was supported by State Research Funding, Hospital District of Southwest Finland, Finnish Cultural, Yrjö Jahnsson and Wihuri foundations, and by Tirkkonen family. JK has been supported by the Academy of Finland grants # 265240, 263278, 308248, 312073 and MO by # 297908.

## Author contributions

JR, JK, NS and NL were responsible for the characterization and recruitment of the Finnish twin pairs and provided the study samples. NP and IK provided the samples from Swedish Twin cohorts. MO and JK provided additional Finnish Twin cohort samples and data. MR performed the neuropathological examination of the brain tissue samples, and MK the sample preparation for RRBS under supervision of RLu. MM performed the bioinformatics analysis of the blood RRBS data under supervision of HL. The sample preparation for scRNA-seq was carried out by MK and TL. The bioinformatics analysis of the scRNA-seq data was performed by JS under supervision of AL and LE. The bioinformatics analysis of the brain RRBS data was carried out by BG under supervision of AL, LE and RLu. The genotyping and pyrosequencing analysis were performed by MK, IK and RLu. Nearly all the authors participated in the processing of the preparation of the manuscript and interpretation of the data. All authors revised and approved the manuscript.

---

**Figure EV1.**
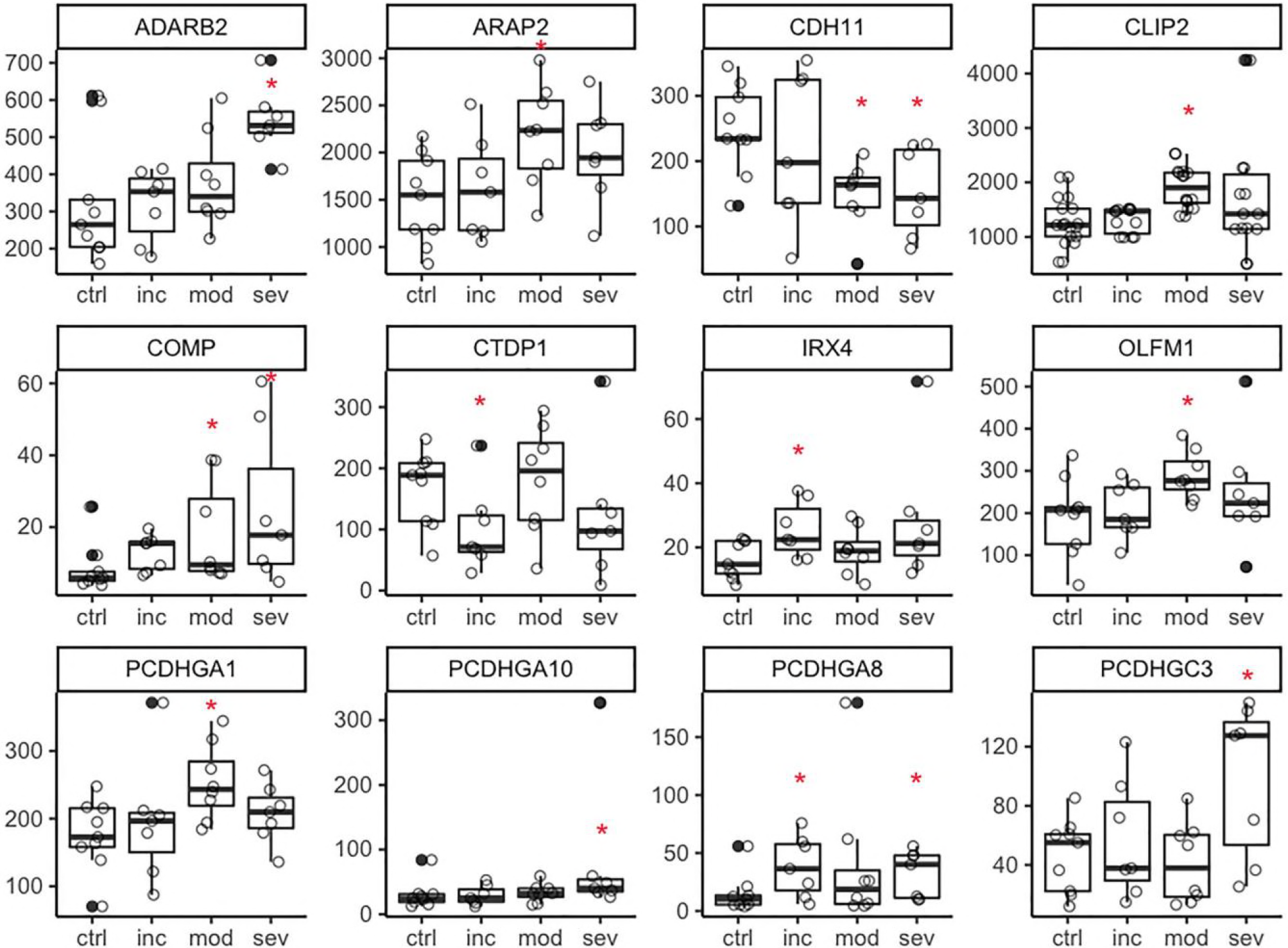
Expression levels of the candidate genes in the data from hippocampus. Expression levels of the genes closest to the AD-associated loci in blood (Table EV1c, Figure 1d) were examined in the data (GDS810) available at GEO, NCBI database. The data included hippocampal samples from control (ctrl, n=9) individuals and individuals with incipient (inc, n=7), moderate (mod, n=8) or severe (sev, n=7) Alzheimer’s disease. The data was analysed with GEO2R with default settings. The disease groups were compared to the control group. The genes with potential expression difference in one of the disease groups (nominal p≤0.05, red asterisk) in comparison to the control group are visualised in the figure (see Table EV2a for details).

**Figure EV2.**
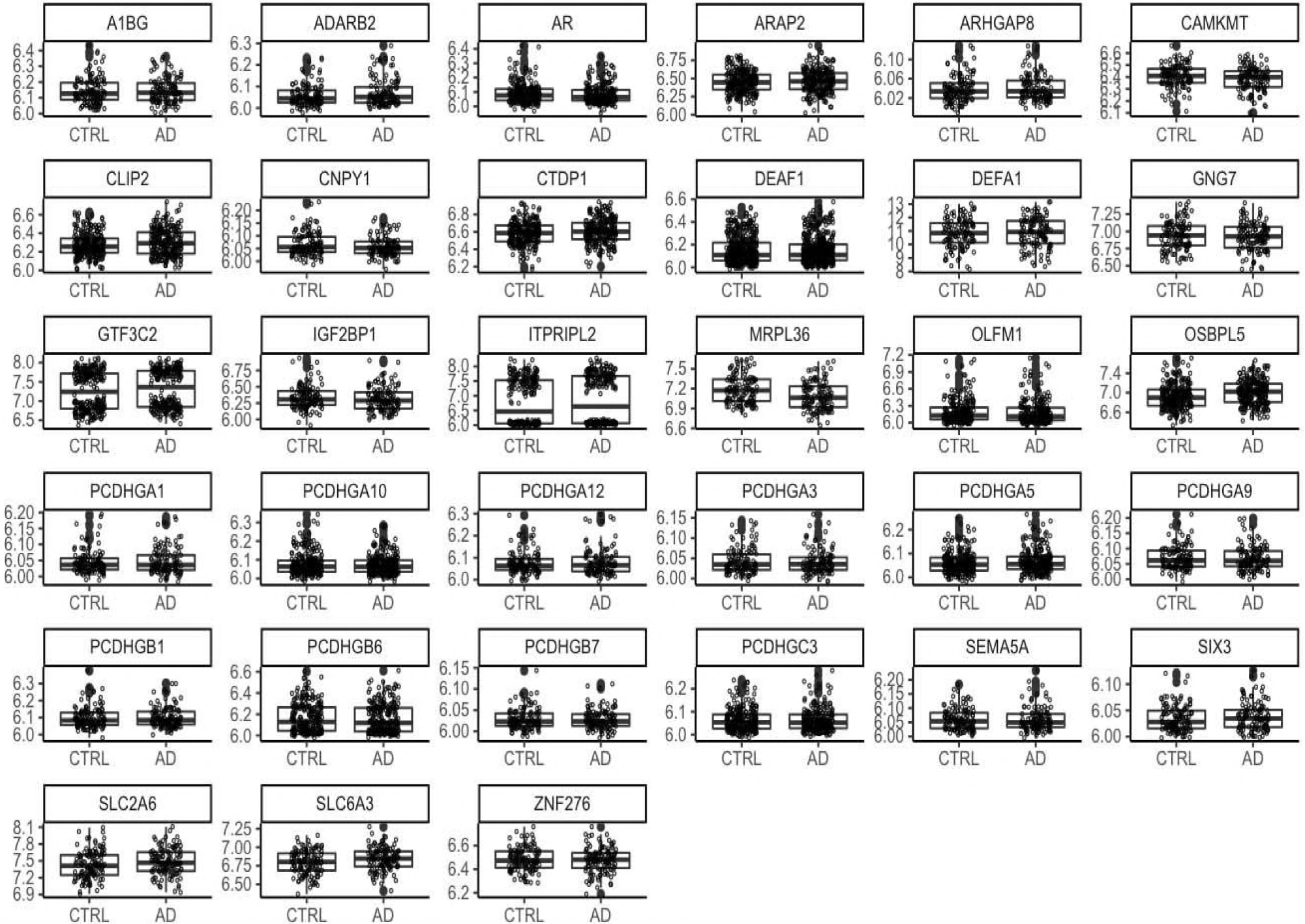
Expression levels of the candidate genes in the data from blood. Expression levels of the genes closest to the AD-associated loci in blood (Table EV1c, Figure 1d) were examined in the data (GSE63061) available at GEO, NCBI database. The data included blood samples from control (CTRL, n=122) individuals and individuals with mild cognitive impairment (n=85) or Alzheimer’s disease (AD, n=121). The data was analysed with GEO2R with default settings. The disease groups were compared to the control group, however, did not differ as the fold changes between the groups were close to 1. The normalised gene expression levels for the control and Alzheimer’s disease groups are visualised in the figure (see Table EV2b for details).

**Figure EV3.**
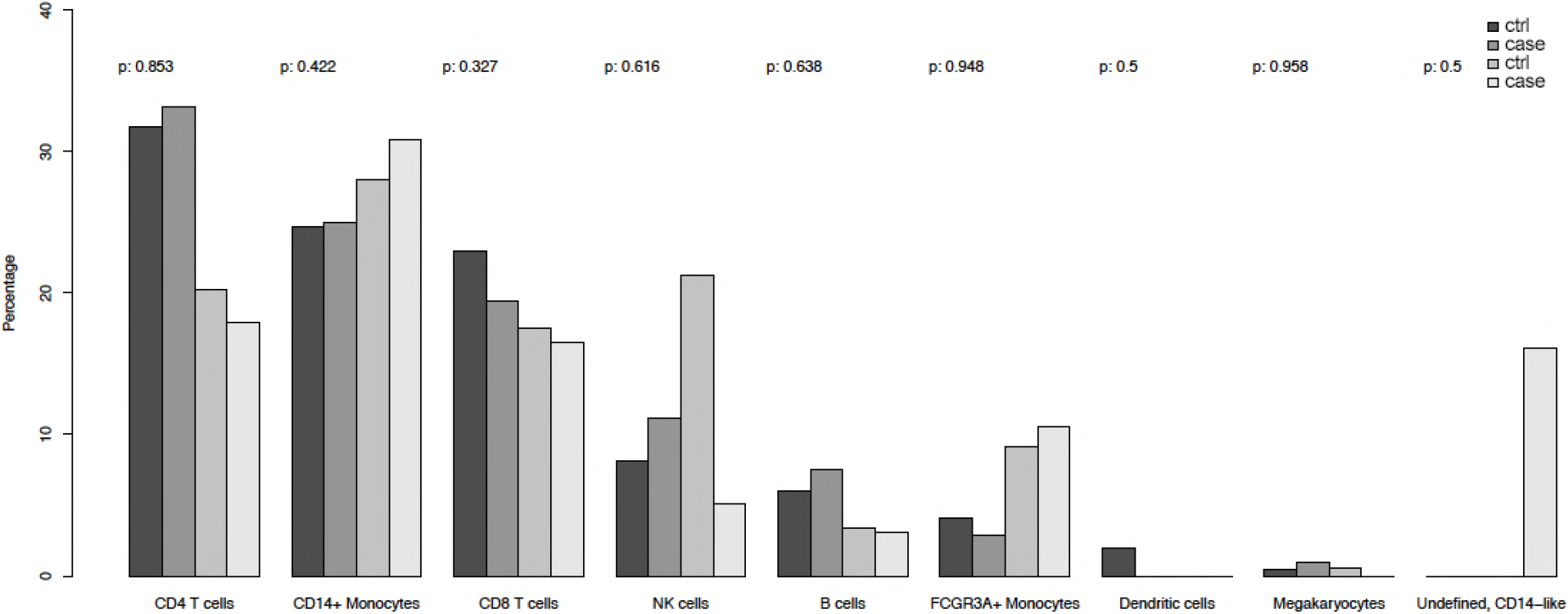
Proportion of peripheral blood mononuclear cell subtypes in twin pairs discordant for episodic memory as determined by single cell 3’ RNA-sequencing.

**Figure EV4. Expression levels of the candidate genes in the peripheral blood mononuclear cell subtypes in twin pairs discordant for episodic memory as determined by single cell 3’ RNA-sequencing.**

**Figure EV5.**
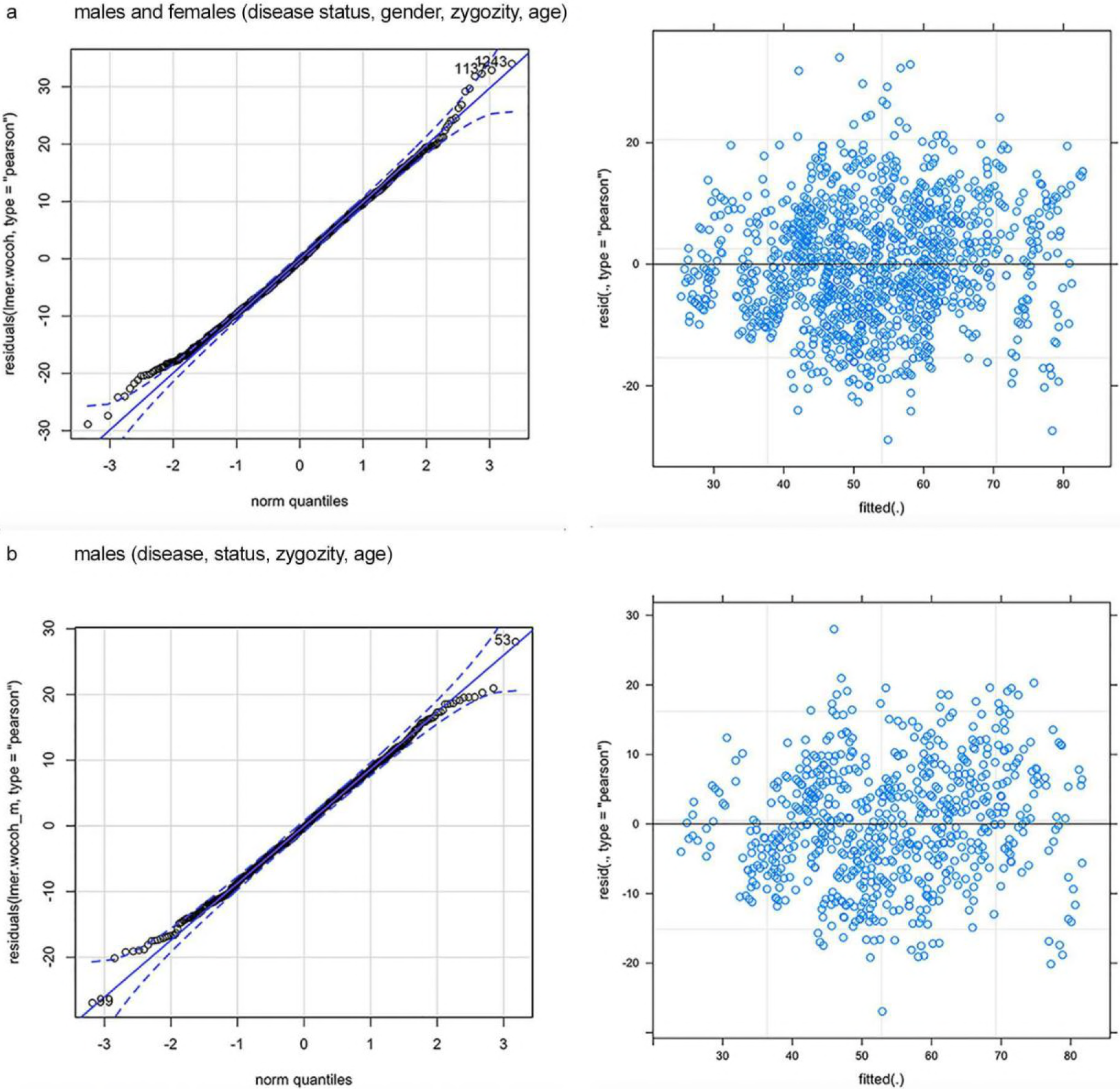
Residual plots for linear mixed effects models. CpG methylation of the region chr10:1,405,405-366, in exon three of *ADARB2* gene was analysed with targeted pyrosequencing. Association of disease status with methylation level (outcome) was examined with linear mixed effects model (lme4 R package) including zygosity, age and gender as fixed effects and twin pair information nested with genomic position as random effects. In the Figure is the representative data of the residual plots for the models: **a**) me ∼ dis + zyg + age + sex + dis * zyg + dis * age + dis * sex + (1 | pairid/pos) in the data including both male and female twin pairs, **b**) me ∼ dis + zyg + age + dis * zyg + dis * age + (1 | pairid/pos) in the data including only males. me=CpG methylation level, dis=disease status, zyg=zygosity, pairid=twin pair information, pos=genomic position.

## References

Akalin A, Kormaksson M, Li S, Garrett-Bakelman FE, Figueroa ME, Melnick A, Mason CE (2012) methylKit: a comprehensive R package for the analysis of genome-wide DNA methylation profiles. Genome Biol 13: R87-2012-13-10-r87.

Alzheimer’s Association (2016) 2016 Alzheimer’s disease facts and figures. Alzheimers Dement 12: 459–509.

Anholt RR (2014) Olfactomedin proteins: central players in development and disease. Front Cell Dev Biol 2: 6.

Barrett T, Wilhite SE, Ledoux P, Evangelista C, Kim IF, Tomashevsky M, Marshall KA, Phillippy KH, Sherman PM, Holko M, Yefanov A, Lee H, Zhang N, Robertson CL, Serova N, Davis S, Soboleva A (2013) NCBI GEO: archive for functional genomics data sets--update. Nucleic Acids Res 41: D991–5.

Bartelt-Kirbach B, Langer-Fischer K, Golenhofen N (2010) Different regulation of N-cadherin and cadherin-11 in rat hippocampus. Cell Commun Adhes 17: 75–82.

Bates D, Mächler M, Bolker B, Walker S (2015) Fitting Linear Mixed-Effects Models Using lme4. Journal of Statistical Software 67: 1–48.

Bertelsen B, Melchior L, Jensen LR, Groth C, Nazaryan L, Debes NM, Skov L, Xie G, Sun W, Brondum-Nielsen K, Kuss AW, Chen W, Tumer Z (2015) A t(3;9)(q25.1;q34.3) translocation leading to OLFM1 fusion transcripts in Gilles de la Tourette syndrome, OCD and ADHD. Psychiatry Res 225: 268–275.

Blalock EM, Geddes JW, Chen KC, Porter NM, Markesbery WR, Landfield PW (2004) Incipient Alzheimer’s disease: microarray correlation analyses reveal major transcriptional and tumor suppressor responses. Proc Natl Acad Sci U S A 101: 2173–2178.

Boyle P, Clement K, Gu H, Smith ZD, Ziller M, Fostel JL, Holmes L, Meldrim J, Kelley F, Gnirke A, Meissner A (2012) Gel-free multiplexed reduced representation bisulfite sequencing for large-scale DNA methylation profiling. Genome Biol 13: R92.

Butler A, Hoffman P, Smibert P, Papalexi E, Satija R (2018) Integrating single-cell transcriptomic data across different conditions, technologies, and species. Nat Biotechnol 36: 411–420.

Cuyvers E, Sleegers K (2016) Genetic variations underlying Alzheimer’s disease: evidence from genome-wide association studies and beyond. Lancet Neurol 15: 857–868.

De Jager PL, Srivastava G, Lunnon K, Burgess J, Schalkwyk LC, Yu L, Eaton ML, Keenan BT, Ernst J, McCabe C, Tang A, Raj T, Replogle J, Brodeur W, Gabriel S, Chai HS, Younkin C, Younkin SG, Zou F, Szyf M, et al (2014) Alzheimer’s disease: early alterations in brain DNA methylation at ANK1, BIN1, RHBDF2 and other loci. Nat Neurosci 17: 1156–1163.

Deming Y, Li Z, Kapoor M, Harari O, Del-Aguila JL, Black K, Carrell D, Cai Y, Fernandez MV, Budde J, Ma S, Saef B, Howells B, Huang KL, Bertelsen S, Fagan AM, Holtzman DM, Morris JC, Kim S, Saykin AJ, et al (2017) Genome-wide association study identifies four novel loci associated with Alzheimer’s endophenotypes and disease modifiers. Acta Neuropathol 133: 839–856.

Dolzhenko E, Smith AD (2014) Using beta-binomial regression for high-precision differential methylation analysis in multifactor whole-genome bisulfite sequencing experiments. BMC Bioinformatics 15: 215-2105-15-215.

Duan Y, Wang SH, Song J, Mironova Y, Ming GL, Kolodkin AL, Giger RJ (2014) Semaphorin 5A inhibits synaptogenesis in early postnatal- and adult-born hippocampal dentate granule cells. Elife 3: 10.7554/eLife.04390.

Dubois B, Hampel H, Feldman HH, Scheltens P, Aisen P, Andrieu S, Bakardjian H, Benali H, Bertram L, Blennow K, Broich K, Cavedo E, Crutch S, Dartigues JF, Duyckaerts C, Epelbaum S, Frisoni GB, Gauthier S, Genthon R, Gouw AA, et al (2016) Preclinical Alzheimer’s disease: Definition, natural history, and diagnostic criteria. Alzheimers Dement 12: 292–323.

Edgar R, Domrachev M, Lash AE (2002) Gene Expression Omnibus: NCBI gene expression and hybridization array data repository. Nucleic Acids Res 30: 207–210.

Fagerberg L, Hallstrom BM, Oksvold P, Kampf C, Djureinovic D, Odeberg J, Habuka M, Tahmasebpoor S, Danielsson A, Edlund K, Asplund A, Sjostedt E, Lundberg E, Szigyarto CA, Skogs M, Takanen JO, Berling H, Tegel H, Mulder J, Nilsson P, et al (2014) Analysis of the human tissue-specific expression by genome-wide integration of transcriptomics and antibody-based proteomics. Mol Cell Proteomics 13: 397–406.

Fanous AH, Zhou B, Aggen SH, Bergen SE, Amdur RL, Duan J, Sanders AR, Shi J, Mowry BJ, Olincy A, Amin F, Cloninger CR, Silverman JM, Buccola NG, Byerley WF, Black DW, Freedman R, Dudbridge F, Holmans PA, Ripke S, et al (2012) Genome-wide association study of clinical dimensions of schizophrenia: polygenic effect on disorganized symptoms. Am J Psychiatry 169: 1309–1317.

Finkel D, Reynolds CA, McArdle JJ, Pedersen NL (2005) The longitudinal relationship between processing speed and cognitive ability: genetic and environmental influences. Behav Genet 35: 535–549.

Fox J, Weisberg S (2011) An R Companion to Applied Regression, Second Edition. Thousand Oaks CA. Sage.

Fransquet PD, Lacaze P, Saffery R, McNeil J, Woods R, Ryan J (2018) Blood DNA methylation as a potential biomarker of dementia: A systematic review. Alzheimers Dement 14: 81–103.

Gaisler-Salomon I, Kravitz E, Feiler Y, Safran M, Biegon A, Amariglio N, Rechavi G (2014) Hippocampus-specific deficiency in RNA editing of GluA2 in Alzheimer’s disease. Neurobiol Aging 35: 1785–1791.

Gatz M, Fratiglioni L, Johansson B, Berg S, Mortimer JA, Reynolds CA, Fiske A, Pedersen NL (2005) Complete ascertainment of dementia in the Swedish Twin Registry: the HARMONY study. Neurobiol Aging 26: 439–447.

Gatz M, Pedersen NL, Berg S, Johansson B, Johansson K, Mortimer JA, Posner SF, Viitanen M, Winblad B, Ahlbom A (1997) Heritability for Alzheimer’s disease: the study of dementia in Swedish twins. J Gerontol A Biol Sci Med Sci 52: M117–25.

Gatz M, Reynolds CA, Fratiglioni L, Johansson B, Mortimer JA, Berg S, Fiske A, Pedersen NL (2006) Role of genes and environments for explaining Alzheimer disease. Arch Gen Psychiatry 63: 168–174.

Goes FS, McGrath J, Avramopoulos D, Wolyniec P, Pirooznia M, Ruczinski I, Nestadt G, Kenny EE, Vacic V, Peters I, Lencz T, Darvasi A, Mulle JG, Warren ST, Pulver AE (2015) Genome-wide association study of schizophrenia in Ashkenazi Jews. Am J Med Genet B Neuropsychiatr Genet 168: 649–659.

Graham N, Arai M, Hagströmer B (2018) multiwayvcov: Multi-Way Standard Error Clustering version 1.2.3 2018.

Gu LZ, Jiang T, Cheng ZH, Zhang YC, Ou MM, Chen MC, Ling WM (2015) TSNARE1 polymorphisms are associated with schizophrenia susceptibility in Han Chinese. J Neural Transm (Vienna) 122: 929–932.

Jarvenpaa T, Laakso MP, Rossi R, Koskenvuo M, Kaprio J, Raiha I, Kurki T, Laine M, Frisoni GB, Rinne JO (2004) Hippocampal MRI volumetry in cognitively discordant monozygotic twin pairs. J Neurol Neurosurg Psychiatry 75: 116–120.

Jarvenpaa T, Raiha I, Kaprio J, Koskenvuo M, Laine M, Kurki T, Vahlberg T, Viljanen T, Ahonen K, Rinne JO (2003) Regional cerebral glucose metabolism in monozygotic twins discordant for Alzheimer’s disease. Dement Geriatr Cogn Disord 16: 245–252.

Jarvenpaa T, Rinne JO, Raiha I, Koskenvuo M, Lopponen M, Hinkka S, Kaprio J (2002) Characteristics of two telephone screens for cognitive impairment. Dement Geriatr Cogn Disord 13: 149–155.

Kaprio J (2013) The Finnish Twin Cohort Study: an update. Twin Res Hum Genet 16: 157–162.

Kaprio J, Koskenvuo M (2002) Genetic and environmental factors in complex diseases: the older Finnish Twin Cohort. Twin Res 5: 358–365.

Konki M, Pasumarthy K, Malonzo M, Sainio A, Valensisi C, Soderstrom M, Emani MR, Stubb A, Narva E, Ghimire B, Laiho A, Jarvelainen H, Lahesmaa R, Lahdesmaki H, Hawkins RD, Lund RJ (2016) Epigenetic Silencing of the Key Antioxidant Enzyme Catalase in Karyotypically Abnormal Human Pluripotent Stem Cells. Sci Rep 6: 22190.

Krueger F (2015) Trim Galore! 2015.

Krueger F, Andrews S (2011) Bismark: a flexible aligner and methylation caller for Bisulfite-Seq applications. Bioinformatics 27: 1571–1572.

Lambert JC, Ibrahim-Verbaas CA, Harold D, Naj AC, Sims R, Bellenguez C, DeStafano AL, Bis JC, Beecham GW, Grenier-Boley B, Russo G, Thorton-Wells TA, Jones N, Smith AV, Chouraki V, Thomas C, Ikram MA, Zelenika D, Vardarajan BN, Kamatani Y, et al (2013) Meta-analysis of 74,046 individuals identifies 11 new susceptibility loci for Alzheimer’s disease. Nat Genet 45: 1452–1458.

Lee E, Giovanello KS, Saykin AJ, Xie F, Kong D, Wang Y, Yang L, Ibrahim JG, Doraiswamy PM, Zhu H (2017) Single-nucleotide polymorphisms are associated with cognitive decline at Alzheimer’s disease conversion within mild cognitive impairment patients. Alzheimers Dement (Amst) 8: 86–95.

Li QS, Parrado AR, Samtani MN, Narayan VA, Alzheimer’s Disease Neuroimaging Initiative (2015) Variations in the FRA10AC1 Fragile Site and 15q21 Are Associated with Cerebrospinal Fluid Abeta1-42 Level. PLoS One 10: e0134000.

Lunnon K, Smith R, Hannon E, De Jager PL, Srivastava G, Volta M, Troakes C, Al-Sarraj S, Burrage J, Macdonald R, Condliffe D, Harries LW, Katsel P, Haroutunian V, Kaminsky Z, Joachim C, Powell J, Lovestone S, Bennett DA, Schalkwyk LC, et al (2014) Methylomic profiling implicates cortical deregulation of ANK1 in Alzheimer’s disease. Nat Neurosci 17: 1164–1170.

Magnusson PK, Almqvist C, Rahman I, Ganna A, Viktorin A, Walum H, Halldner L, Lundstrom S, Ullen F, Langstrom N, Larsson H, Nyman A, Gumpert CH, Rastam M, Anckarsater H, Cnattingius S, Johannesson M, Ingelsson E, Klareskog L, de Faire U, et al (2013) The Swedish Twin Registry: establishment of a biobank and other recent developments. Twin Res Hum Genet 16: 317–329.

Mladenova D, Barry G, Konen LM, Pineda SS, Guennewig B, Avesson L, Zinn R, Schonrock N, Bitar M, Jonkhout N, Crumlish L, Kaczorowski DC, Gong A, Pinese M, Franco GR, Walkley CR, Vissel B, Mattick JS (2018) Adar3 Is Involved in Learning and Memory in Mice. Front Neurosci 12: 243.

Nakaya N, Sultana A, Lee HS, Tomarev SI (2012) Olfactomedin 1 interacts with the Nogo A receptor complex to regulate axon growth. J Biol Chem 287: 37171–37184.

Nakaya N, Sultana A, Munasinghe J, Cheng A, Mattson MP, Tomarev SI (2013) Deletion in the N-terminal half of olfactomedin 1 modifies its interaction with synaptic proteins and causes brain dystrophy and abnormal behavior in mice. Exp Neurol 250: 205–218.

Nakazawa M (2017) fmsb: Functions for Medical Statistics Book with some Demographic Data version 0.6.1 2018.

Nordberg A, Rinne JO, Kadir A, Langstrom B (2010) The use of PET in Alzheimer disease. Nat Rev Neurol 6: 78–87.

Oakes E, Anderson A, Cohen-Gadol A, Hundley HA (2017) Adenosine Deaminase That Acts on RNA 3 (ADAR3) Binding to Glutamate Receptor Subunit B Pre-mRNA Inhibits RNA Editing in Glioblastoma. J Biol Chem 292: 4326–4335.

Querfurth HW, LaFerla FM (2010) Alzheimer’s disease. N Engl J Med 362: 329–344.

R Core Team (2017) R: A language and environment for statistical computing. R Foundation for Statistical Computing, Vienna, Austria. 2017–2018.

Reitz C, Brayne C, Mayeux R (2011) Epidemiology of Alzheimer disease. Nat Rev Neurol 7: 137–152.

Reitz C, Tosto G, Vardarajan B, Rogaeva E, Ghani M, Rogers RS, Conrad C, Haines JL, Pericak-Vance MA, Fallin MD, Foroud T, Farrer LA, Schellenberg GD, George-Hyslop PS, Mayeux R, Alzheimer’s Disease Genetics Consortium (ADGC) (2013) Independent and epistatic effects of variants in VPS10-d receptors on Alzheimer disease risk and processing of the amyloid precursor protein (APP). Transl Psychiatry 3: e256.

Roubroeks JAY, Smith RG, van den Hove DLA, Lunnon K (2017) Epigenetics and DNA methylomic profiling in Alzheimer’s disease and other neurodegenerative diseases. J Neurochem.

Scheinin NM, Aalto S, Kaprio J, Koskenvuo M, Raiha I, Rokka J, Hinkka-Yli-Salomaki S, Rinne JO (2011) Early detection of Alzheimer disease: (1)(1)C-PiB PET in twins discordant for cognitive impairment. Neurology 77: 453–460.

Scheinin NM, Scheinin M, Rinne JO (2011) Amyloid imaging as a surrogate marker in clinical trials in Alzheimer’s disease. Q J Nucl Med Mol Imaging 55: 265–279.

Sleiman P, Wang D, Glessner J, Hadley D, Gur RE, Cohen N, Li Q, Hakonarson H, Janssen-CHOP Neuropsychiatric Genomics Working Group (2013) GWAS meta analysis identifies TSNARE1 as a novel Schizophrenia / Bipolar susceptibility locus. Sci Rep 3: 3075.

Sood S, Gallagher IJ, Lunnon K, Rullman E, Keohane A, Crossland H, Phillips BE, Cederholm T, Jensen T, van Loon LJ, Lannfelt L, Kraus WE, Atherton PJ, Howard R, Gustafsson T, Hodges A, Timmons JA (2015) A novel multi-tissue RNA diagnostic of healthy ageing relates to cognitive health status. Genome Biol 16: 185-015-0750-x.

Tan MH, Li Q, Shanmugam R, Piskol R, Kohler J, Young AN, Liu KI, Zhang R, Ramaswami G, Ariyoshi K, Gupte A, Keegan LP, George CX, Ramu A, Huang N, Pollina EA, Leeman DS, Rustighi A, Goh YPS, GTEx Consortium, et al (2017) Dynamic landscape and regulation of RNA editing in mammals. Nature 550: 249–254.

Therneau T (2018a) coxme: Mixed Effects Cox Models. https://CRAN.Rproject.org/package=coxme. 2018.

Therneau T (2018b) A Package for Survival Analysis in S. version 2.38, https://CRAN.Rproject.org/package=survival. 2018.

Tsai PC, Bell JT (2015) Power and sample size estimation for epigenome-wide association scans to detect differential DNA methylation. Int J Epidemiol 44: 1429–1441.

Virta JJ, Karrasch M, Kaprio J, Koskenvuo M, Raiha I, Viljanen T, Rinne JO (2008) Cerebral glucose metabolism in dizygotic twin pairs discordant for Alzheimer’s disease. Dement Geriatr Cogn Disord 25: 9–16.

Vulto-van Silfhout AT, Rajamanickam S, Jensik PJ, Vergult S, de Rocker N, Newhall KJ, Raghavan R, Reardon SN, Jarrett K, McIntyre T, Bulinski J, Ownby SL, Huggenvik JI, McKnight GS, Rose GM, Cai X, Willaert A, Zweier C, Endele S, de Ligt J, et al (2014) Mutations affecting the SAND domain of DEAF1 cause intellectual disability with severe speech impairment and behavioral problems. Am J Hum Genet 94: 649–661.

Watson CT, Roussos P, Garg P, Ho DJ, Azam N, Katsel PL, Haroutunian V, Sharp AJ (2016) Genome-wide DNA methylation profiling in the superior temporal gyrus reveals epigenetic signatures associated with Alzheimer’s disease. Genome Med 8: 5-015-0258-8.

Wickham H (2009) ggplot2: Elegant Graphics for Data Analysis. Springer-Verlag New York.

Zeileis A, Hothorn T (2002) Diagnostic Checking in Regression Relationships. R News 2: 7–10.

Zhao J, Zhu Y, Yang J, Li L, Wu H, De Jager PL, Jin P, Bennett DA (2017) A genome-wide profiling of brain DNA hydroxymethylation in Alzheimer’s disease. Alzheimers Dement 13: 674–688.

